# STN-DBS frequency-tuned beta subband dynamics are associated with speech tempo in Parkinson’s disease

**DOI:** 10.64898/2026.06.07.729490

**Authors:** Haiyu Tian, Yuxuan Wu, Yu Fan, Zixiao Yin, Yi Guo, Fangang Meng, Jianguo Zhang, Huiling Yu, Luming Li

## Abstract

In Parkinson’s disease, subthalamic nucleus deep brain stimulation (STN-DBS) frequency modulates narrow beta subbands, with low-frequency stimulation (LFS) selectively attenuating 15–23 Hz power. The negative association between this subband and speech tempo, together with the LFS-related improvement in articulatory rate, suggests that 15–23 Hz subband dynamics may serve as a candidate neural marker associated with speech tempo modulation.

## Main text

Stimulation frequency is a key programming parameter in deep brain stimulation (DBS). In subthalamic nucleus DBS (STN-DBS) for Parkinson’s disease (PD), high frequency stimulation (HFS) has been widely shown to improve cardinal motor symptoms including bradykinesia, tremor, and rigidity ^1,2^. Nevertheless, in some cases, motor benefits may be accompanied by side effects on speech function^2,3^. Low frequency stimulation (LFS) has been reported to improve speech^4-7^. However, existing findings remain inconsistent^3^. Most previous studies have largely relied on behavioral observations, with limited evidence linking neurophysiological and behavioral outcomes.

The brain can be conceptualized as a state-dependent dynamical system shaped by multiscale endogenous neural oscillations^8,9^. The frequency of external electrical stimulation may modulate intrinsic oscillatory activity, thereby influencing neurophysiological processes and behavioral performance^4,10-12^. In PD, exaggerated beta band (13–35 Hz)^13^ oscillations in the STN are associated with motor symptoms such as bradykinesia and rigidity^13,14^, and represent a typical biomarker for DBS modulation. However, despite evidence that STN-DBS frequency modulates beta oscillations^15,16^, the behavioral relevance of frequency-specific beta dynamics for speech and motor function remains unclear. Previous work using constant-voltage stimulation reported that 60 Hz STN-DBS selectively amplified oscillatory activity in the 10–15 Hz range compared with 140 Hz stimulation, without worsening bradykinesia^16^. This suggests that stimulation frequency may regulate STN oscillations in narrow frequency-specific bands. However, changing the stimulation frequency under constant-amplitude conditions inevitably alters the total electrical energy delivered (TEED). Such TEED changes may affect speech performance via motor function changes^17^, thereby confounding the effect of stimulation frequency.

In this study, we used sensing-enabled STN-DBS devices^18^ to deliver stimulation at 60, 90, 130, and 180 Hz under approximately constant TEED, while simultaneously recording subthalamic local field potentials (STN-LFPs), assessing MDS-UPDRS Part III motor scores and speech features from multiple tasks under both off- and on-medication conditions. We aimed to evaluate how STN-DBS stimulation frequency affects beta band oscillatory activity, speech, and motor performance, and to further examine the associations between LFP power and speech features, as well as between speech and motor function.

Under approximately constant TEED, stimulation frequency did not induce significant changes in power within the canonical low beta (13–20 Hz) or high beta (20–35 Hz) bands. We screened all beta-band subbands with integer-Hz boundaries and bandwidths of ≥1 Hz, and found that LFS (60 Hz) produced significantly greater suppression of the 15–23 Hz subband than HFS (180 Hz) in the on-medication condition (Fig. 1d, *p = 0*.*011*), with a consistent trend observed off-medication (Supplementary Fig. 1a), suggesting preferential attenuation of this lower-beta subband by LFS across medication states. In contrast, HFS showed greater suppression than LFS only in the higher 29–35 Hz subband, and only in the off-medication condition (Fig. 1c, *p = 0*.*021*).

**Fig. 1.**
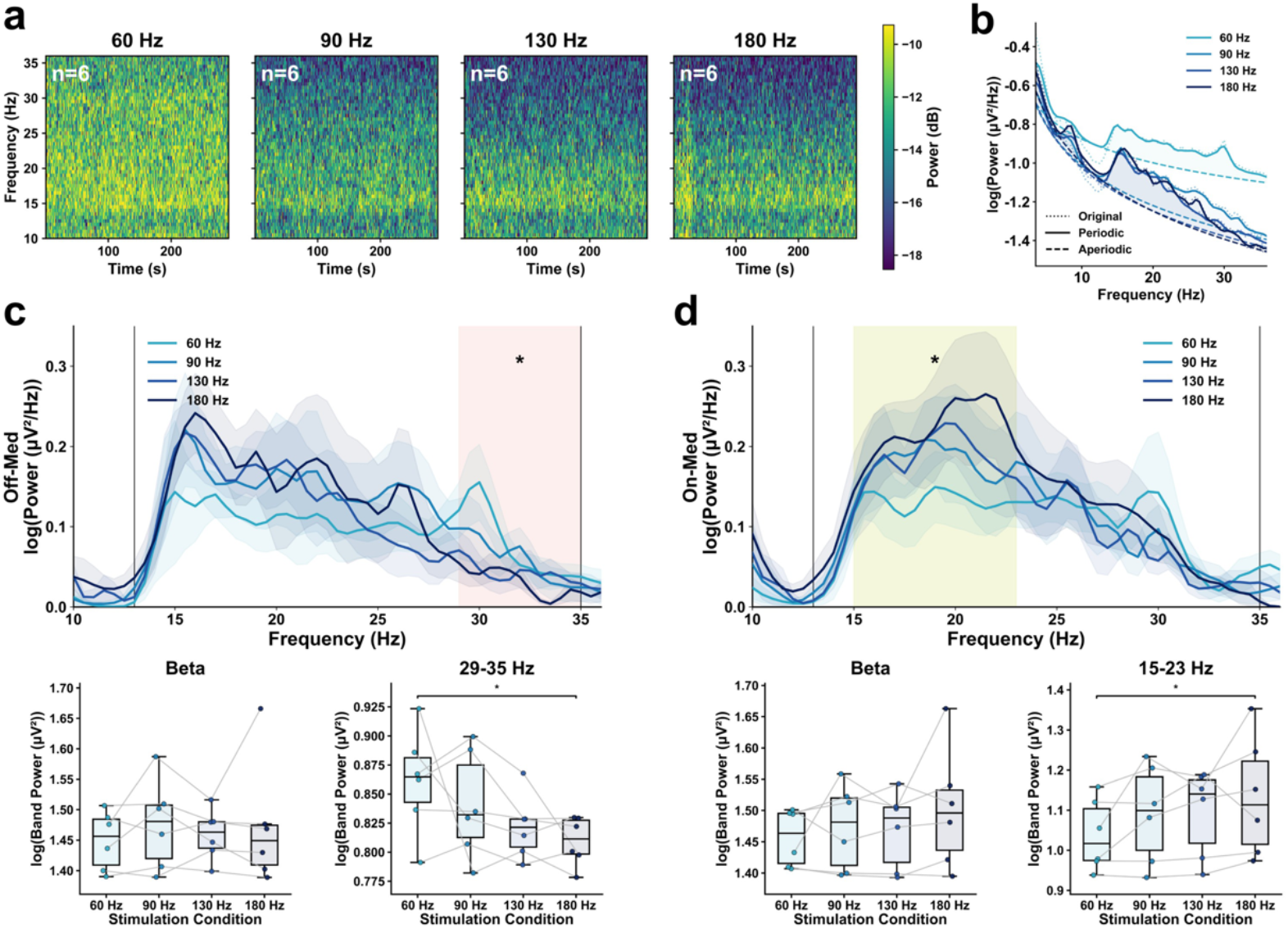
STN-LFP oscillatory activity under different DBS frequencies. **a** Power spectral characteristics of STN-LFPs across four stimulation frequencies (60, 90, 130, and 180 Hz, off-medication condition, before FOOOF). **b** FOOOF-based decomposition of simultaneously recorded STN-LFP power spectra into periodic and aperiodic components. **c** STN-LFP oscillatory activity under different stimulation frequencies in the off-medication condition. **d** STN-LFP oscillatory activity under different stimulation frequencies in the on-medication condition.

We next investigated the effects of stimulation frequency on speech features. To isolate the frequency-specific effects on speech, in this part we analyzed only the off-medication data. Speech features were normalized within each participant by dividing values obtained under each stimulation condition by the corresponding participant-specific baseline measured under the off-DBS and off-medication baseline. MDS-UPDRS III motor scores were comparable across stimulation frequencies under approximately constant TEED. Under this condition, LFS (60 Hz) significantly improved the diadochokinetic (DDK) rate relative to baseline (Fig. 2b, *p = 0*.*031*). However, direct comparisons across the four stimulation frequencies showed no significant differences in speech features.

**Fig. 2.**
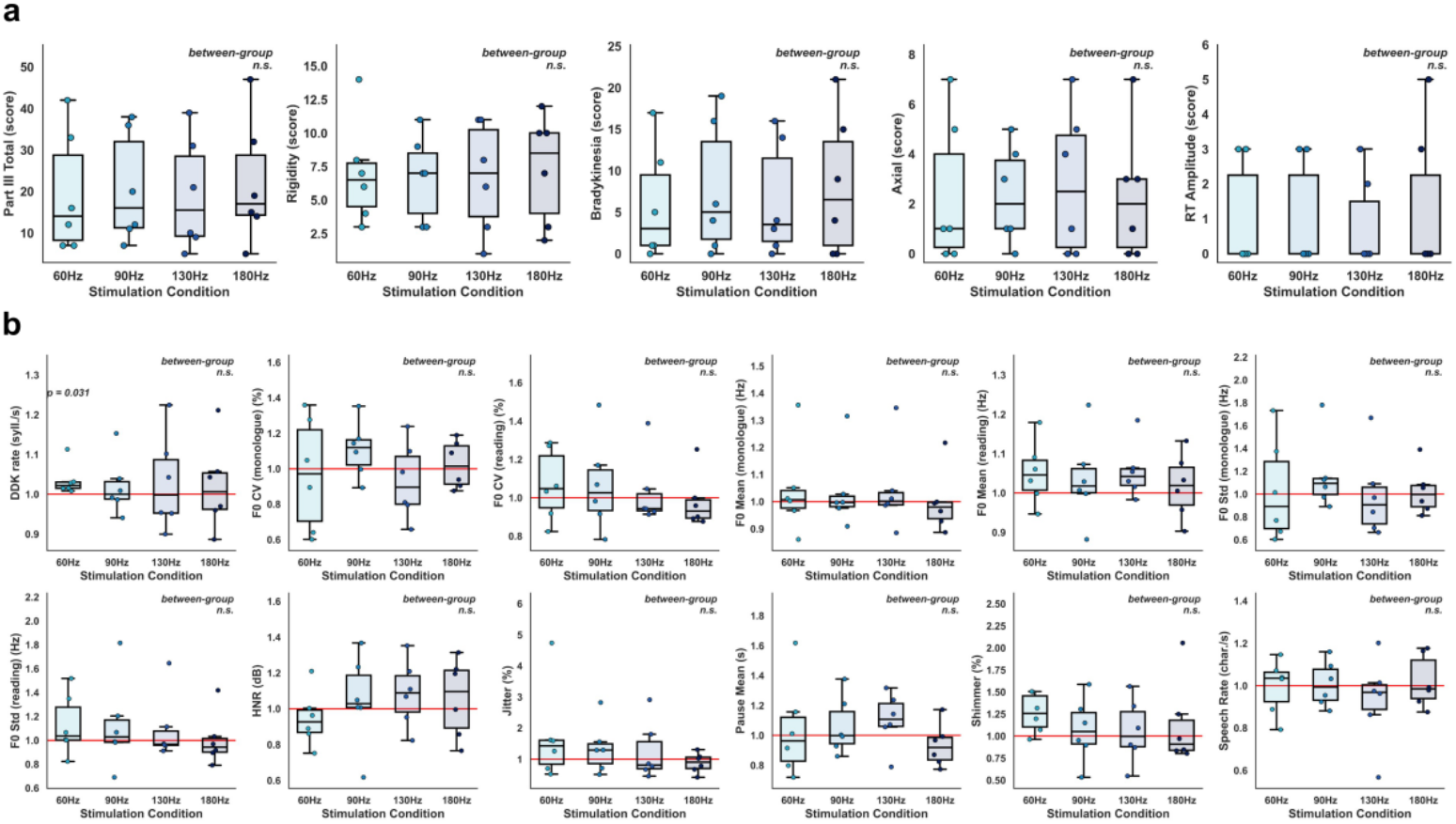
Effects of STN-DBS frequency on MDS-UPDRS Part III motor scores and speech features (in the off-medication condition). **a** MDS-UPDRS Part III motor scores under different stimulation conditions. **b** Speech features under different STN-DBS frequency conditions.

We further examined the associations between normalized band power and speech features, as well as speech features and motor scores. We found that power in the frequency-specific 15–23 Hz subband, canonical beta band, and high beta band were negatively associated with monologue speech rate (Fig. 3a, *β*_15–23 *Hz*_ *= -0*.*258, p*_15–23 *Hz*_ *= 0*.*012*; *β*_*beta*_ *= -0*.*328, p*_*beta*_ *= 0*.*025*; *β*_*highbeta*_ *= -0*.*243, p*_*highbeta*_ *= 0*.*014*). In addition, fundamental frequency coefficient of variation (F0 CV) variability was negatively associated with the MDS-UPDRS III rigidity score, suggesting that reduced F0 CV may be related to greater rigidity severity (Fig. 3b, *β*_*F*0 *CV*(*monologue*)_ *= -0*.*380, p*_*F*0 *CV*(*monologue*)_ *= 0*.*016*; *β*_*F*0 *CV*(*reading*)_ *= -0*.*382, p*_*F*0 *CV*(*reading*)_ *= 0*.*016*). These associations showed broadly similar directions at the group and individual levels. Nevertheless, inter-individual differences were evident in subject-specific trends, feature ranges, and data dispersion, highlighting the importance of accounting for individual response patterns in electrophysiological and behavioral analyses of neuromodulation.

**Fig. 3.**
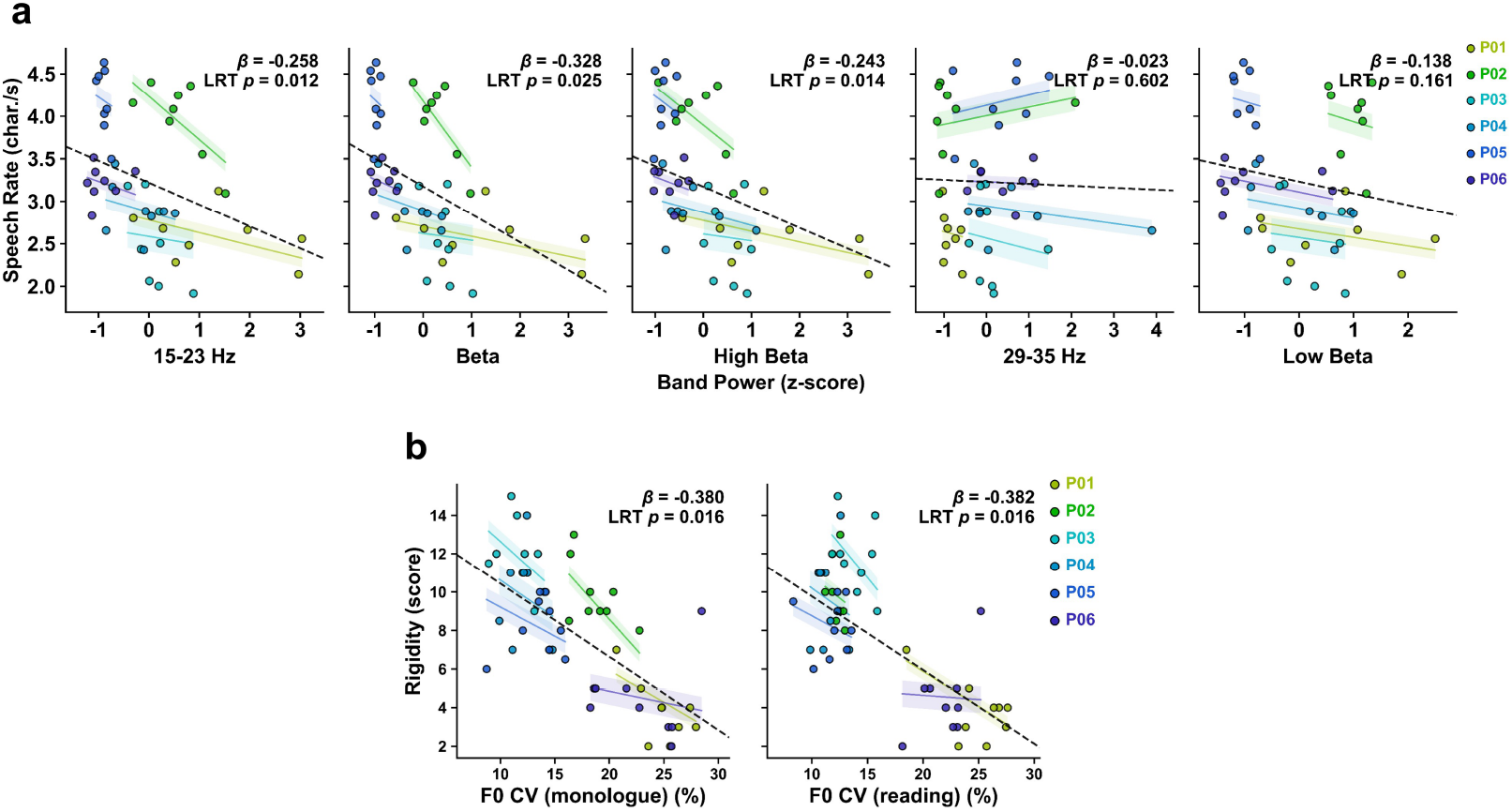
Cross-modal feature associations. **a** Standardized beta band and subband power were negatively associated with monologue speech rate. **b** Fundamental frequency coefficient of variation (F0 CV) was negatively associated with the MDS-UPDRS Part III rigidity score.

Previous studies suggest that low beta oscillations have been more closely linked to dopamine-depleted states and pathological synchronization, whereas high beta oscillations may reflect activity within the cortico-STN hyperdirect pathway and broader motor-network dynamics^13,19,20^. Consistent with this subband-specific view, we identified two frequency-specific oscillatory patterns: LFS (60 Hz) suppressed 15–23 Hz subband activity in the on-medication condition, whereas HFS (180 Hz) suppressed 29–35 Hz subband activity in the off-medication condition. However, effects were not fully consistent across medication states: LFS-related suppression of 15–23 Hz activity showed a similar directional trend in the off-medication condition, whereas HFS-related suppression of 29–35 Hz subband activity was attenuated in the on-medication condition. As 29–35 Hz subband activity falls within the upper high-beta range and may reflect both motor-circuit dynamics and Parkinsonian pathophysiology^13^, the frequency-dependent modulation of this narrow subband should be interpreted cautiously.

Under approximately constant TEED, MDS-UPDRS Part III motor scores did not differ significantly across stimulation frequencies, suggesting that frequency-related speech effects were unlikely to be secondary to global motor changes. Notably, DDK rate significantly increased under LFS, indicating an improved capacity for rapid and repetitive articulatory movements^4^. However, speech rate did not change significantly. Given the greater linguistic and articulatory complexity of continuous speech relative to the DDK task, we cautiously suggest that larger samples and longer monologue recordings are needed to determine whether LFS can yield measurable improvements in natural speech rate. In addition, LFS also showed a nonsignificant trend toward reduced HNR and increased shimmer, suggesting that its effects may be more specific to rhythmic or articulatory-temporal aspects of speech, rather than uniformly improving voice quality.

Cross-modal analyses further showed that monologue speech rate was negatively associated with oscillatory power in the frequency-specific 15–23 Hz subband, canonical beta band, and high-beta band. Increased beta activity may reflect a stronger inhibitory state within the STN–basal ganglia–cortical motor circuit, potentially constraining speech initiation and continuous speech production. In parallel, previous reports have shown that normal speech is accompanied by reductions in STN alpha and beta power, whereas speech-initiation blocks are associated with increased beta activity, particularly within the 15–25 Hz subband^21,22^. Similarly, we found that LFS significantly attenuated 15–23 Hz subband activity, and that power in this narrow subband was negatively associated with speech rate. These findings suggest that LFS-related attenuation of 15–23 Hz activity may represent a candidate neural marker associated with speech tempo modulation. We also observed a negative association between F0 variability and the rigidity score, supporting a link between rigidity-related phonatory impairment and reduced pitch modulation^4^.

This study is limited by its small sample size, short-term design, and fixed stimulation order, which may have influenced the group-level comparisons in the on-medication condition. To address this, we compared power spectral densities (PSDs) between the off-and on-medication conditions and observed low-beta suppression after medication, supporting an effective dopaminergic medication response (Supplementary Fig. 2). Future studies with larger cohorts and longitudinal follow-up are needed to further validate the neural mechanisms underlying DBS frequency-dependent effects and to optimize stimulation protocols for improving speech in PD.

## Methods

### Participants

Six patients with PD were included in this study. All participants underwent bilateral STN-DBS electrode implantation (L301C; Beijing PINS Medical Co., Ltd., Beijing, China), and were connected to a DBS device capable of wireless LFP data acquisition (G106RS, Beijing PINS Medical Co., Ltd.). Electrode reconstruction was performed using the Lead-DBS tool^23^ (Supplementary Fig. 3). The study was approved by the ethics committees of both Beijing Tiantan Hospital and Peking Union Medical College Hospital. Written informed consent was obtained from all participants. The demographic and clinical characteristics are summarized in Table 1.

**Table 1.** Demographic and clinical characteristics.

| **Demographic and Clinical Characteristics** |  |
| --- | --- |
| Number of patients | 6 |
| Age (years) | 60.83 ± 5.23 |
| Gender (Female/Male) | 3/3 |
| PD duration at surgery (years) | 8.50 ± 1.38 |
| Time from DBS surgery to experiment (months) | 9.33 ± 6.35 |
| LED* at the experiment (mg) | 179.75 ± 45.12 |

\*LED, levodopa equivalent dose.

### Experimental procedure

All participants were tested under four DBS frequency conditions in a fixed order: 130 Hz (the initial baseline stimulation frequency; except for one participant who received 140 Hz for therapeutic reasons), 60, 90, and 180 Hz. Testing was first conducted in the off-medication condition after withdrawal of antiparkinsonian medication for at least 12 h and was then repeated in the on-medication condition.

DBS parameters were adjusted to maintain approximately constant TEED. TEED was calculated as follows:

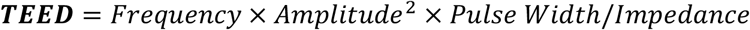

Active contacts and pulse width were controlled across conditions. Amplitude values were adjusted to the nearest 0.05 mA and reported to two decimal places. Normalized TEED across stimulation frequencies was calculated using the baseline stimulation frequency condition as the reference. The results are shown in Table 2, and detailed DBS parameters are provided in Supplementary Table 1.

**Table 2.** Normalized TEED across stimulation frequencies.

| Frequency (Hz) | Hemisphere | Normalized TEED (a.u.) |
| --- | --- | --- |
| 130 Hz | Left-STN | 1 |
|  | Right-STN | 1 |
| 60 Hz | Left-STN | 0.992 ± 0.005 |
|  | Right-STN | 1.004 ± 0.009 |
| 90 Hz | Left-STN | 0.987 ± 0.011 |
|  | Right-STN | 1.000 ± 0.007 |
| 180 Hz | Left-STN | 0.992 ± 0.012 |
|  | Right-STN | 1.001 ± 0.015 |

During each DBS condition, a 10-min wash-in period was applied, followed by 5 min of resting-state STN-LFP recording in simultaneous stimulation-and-recording mode. Multi-task speech assessment and MDS-UPDRS Part III evaluation were then performed.

### LFP recording and preprocessing

LFP data were transmitted in real time, displayed, and stored wirelessly via the sensing-enabled DBS device (G106RS, PINS Medical Co). For each participant, we analyzed the hemisphere with higher signal quality and fewer artifacts. LFP preprocessing was performed in MATLAB R2025a. Signals were bandpass-filtered between 1 and 55 Hz, followed by ECG artifacts removal using a template-based EAS method with 30-s windows. Short-time Fourier transform (STFT) was performed with a 2-s Hanning window and 50% overlap. The FOOOF algorithm^24^ was then used to decompose the simultaneously recorded LFP signals into periodic and aperiodic components.

### Speech features and MDS-UPDRS motor assessments

Speech data were acquired using a microphone (Razer, Singapore) in 44.1 kHz stereo mode and analyzed in Praat (v6.4.23; University of Amsterdam, the Netherlands). Speech recordings were performed in a quiet room, with participants seated comfortably, and the microphone placed on a horizontal tabletop approximately 30 cm from the participant’s mouth. Participants completed four speech tasks in each condition: (1) Sustained phonation of the vowel /a/ in a single breath; (2) The DDK task, in which participants repeatedly produced the syllable sequence /pa-ta-ka/ in a single breath; (3) Reading aloud a 106-character Chinese passage in a comfortable speaking style, which was designed to cover a wide range of Mandarin initials and finals with an approximately balanced distribution of lexical tones; (4) One-minute free-speech task on any topic in a natural speaking style, without reciting, chanting, or singing, which was intended to approximate natural daily communication. The following speech features were extracted from the speech samples for analysis: HNR, Jitter, Shimmer, DDK rate, F0 Mean, F0 Std, F0 CV, Speech Rate. Definitions of the speech tasks and speech features are summarized in Supplementary Table 2.

In addition, the off-DBS and off-medication condition data were obtained from the clinical visit in the same month as the experimental session, using the identical speech testing protocol. Speech outcomes under the 130 Hz and off-medication condition did not differ significantly between the experimental session and the clinical visit (Supplementary Fig. 4), supporting the use of dual-off condition outcomes in this analysis. The motor outcome were the changes in MDS-UPDRS Part III and included: (1) total score; (2) rigidity (3.3); (3) bradykinesia (3.4–3.8); (4) axial (3.9–3.14); (5) resting tremor amplitude (RT amplitude, 3.17).

### Statistical analysis

Statistical analyses were performed in Python 3.12.2. Between-condition effects were tested using linear mixed-effects models (LMMs) with stimulation frequency as a categorical fixed effect and subject as a random intercept, followed by likelihood-ratio tests against the corresponding null models. Multiple comparisons were controlled using the Benjamini–Hochberg false discovery rate procedure. Baseline-normalized speech features were compared with baseline using Wilcoxon signed-rank tests. Cross-modal associations were assessed using LMMs with subject-specific random intercepts and slopes; LFP band power was standardized before model fitting to facilitate comparison of effect trends across frequency bands. Significance was defined as a corrected p < 0.05 where applicable.

## Acknowledgments

The authors thank Lingxiao Guan, Guokun Zhang and Yingtian Liu from the National Engineering Research Center of Neuromodulation, School of Aerospace Engineering, Tsinghua University, for their assistance in data acquisition.

