## Supplementary Figures 1-4; Supplementary Tables 1-2 for "STN-DBS frequency-tuned beta subband dynamics are associated with speech tempo in Parkinson’s disease"

**Supplementary Figure 1.** ***15–23 Hz activity across stimulation frequencies in the off-medication condition (panel a) and its association with DDK rate across medication conditions (panel b).***

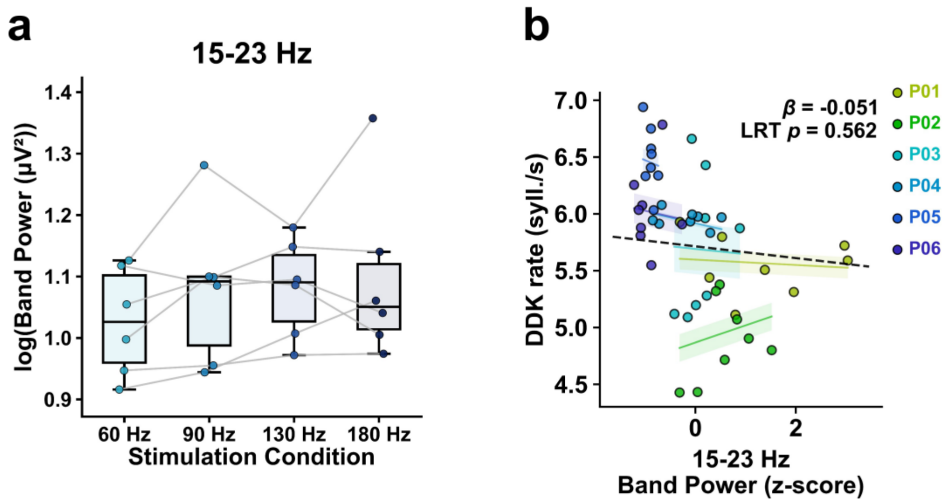

**Supplementary Figure 2.** ***Off- versus on-medication PSD comparisons at each stimulation frequency reveal low beta suppression.***

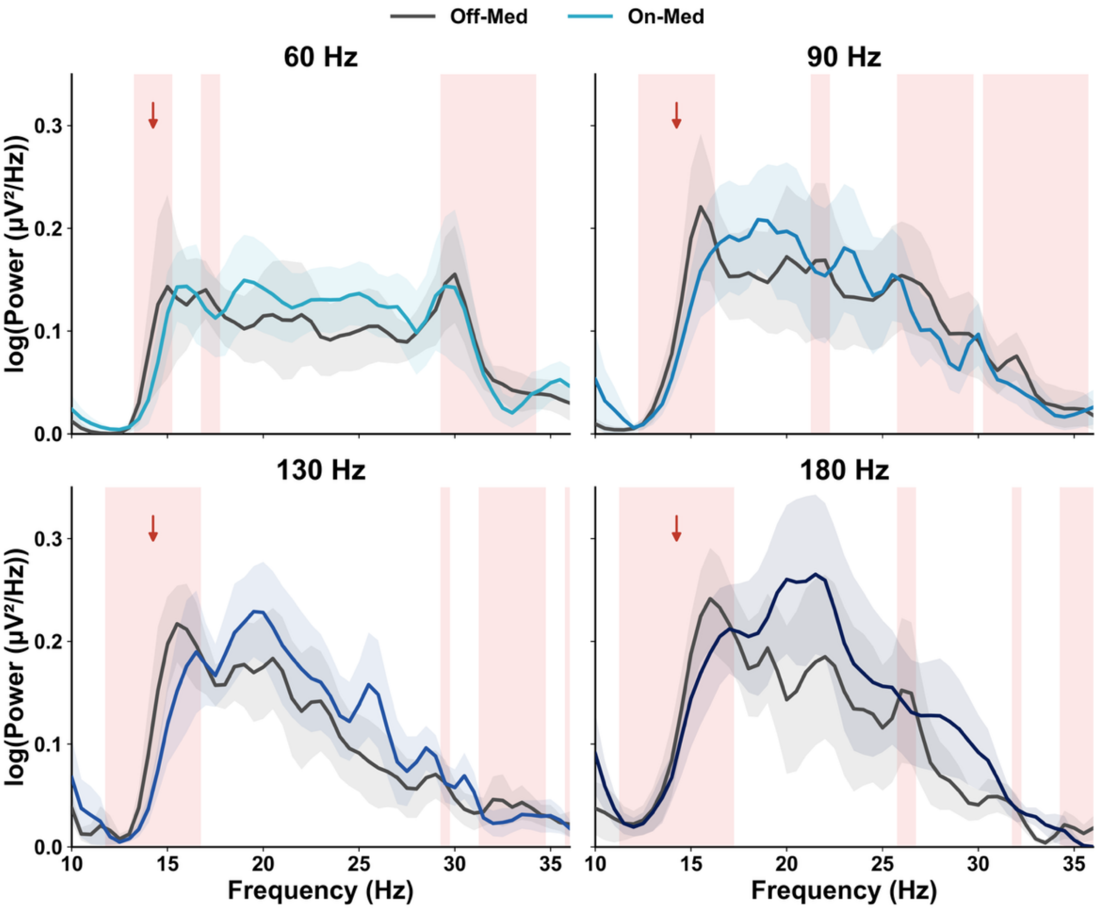

**Supplementary Figure 3.** ***Electrode reconstructions.***

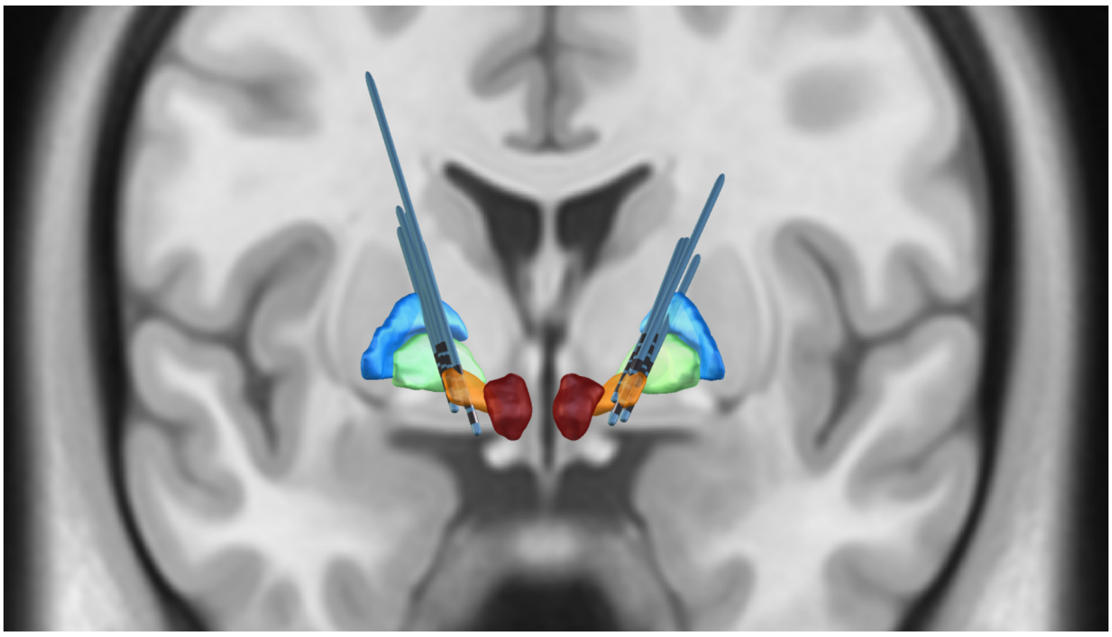

**Supplementary Figure 4.** ***Comparison of speech outcomes under 130-Hz stimulation in the off-medication condition between the frequency-modulation experiment and the follow-up visit (Wilcoxon signed-rank tests).***

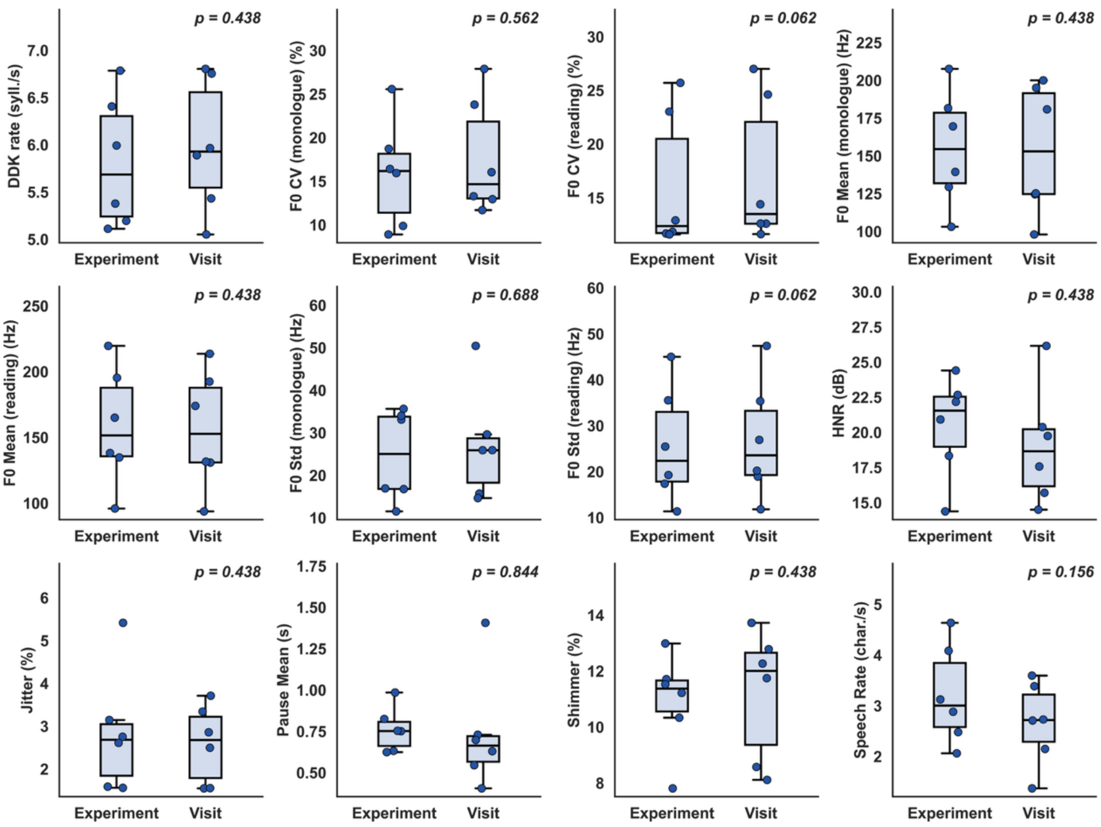

**Supplementary Table 1. DBS parameters and normalized TEED across stimulation conditions.**

| Participant | Hemisphere | Frequency (Hz) | Amplitude (mA) | Pulse width (μs) | Contact | Normalized TEED (a.u.) |
|---|---|---|---|---|---|---|
| P01 | Left-STN | 130 | 1.85 | 70 | C+6- | 1.000 |
| | | 60 | 2.7 | | | 0.983 |
| | | 90 | 2.2 | | | 0.979 |
| | | 180 | 1.55 | | | 0.972 |
| | Right-STN | 130 | 2 | | C+2- | 1.000 |
| | | 60 | 2.95 | | | 1.004 |
| | | 90 | 2.4 | | | 0.997 |
| | | 180 | 1.7 | | | 1.000 |
| P02 | Left-STN | 130 | 2.35 | 70 | C+2- | 1.000 |
| | | 60 | 3.45 | | | 0.995 |
| | | 90 | 2.8 | | | 0.983 |
| | | 180 | 2 | | | 1.003 |
| | Right-STN | 130 | 2.95 | 80 | C+5- | 1.000 |
| | | 60 | 4.35 | | | 1.004 |
| | | 90 | 3.55 | | | 1.003 |
| | | 180 | 2.5 | | | 0.994 |
| P03 | Left-STN | 130 | 2.7 | 70 | C+2- | 1.000 |
| | | 60 | 3.95 | | | 0.988 |
| | | 90 | 3.25 | | | 1.003 |
| | | 180 | 2.3 | | | 1.005 |
| | Right-STN | 130 | 2 | | C+7- | 1.000 |
| | | 60 | 2.95 | | | 1.004 |
| | | 90 | 2.4 | | | 0.997 |
| | | 180 | 1.7 | | | 1.000 |
| P04 | Left-STN | 140 | 2.1 | 80 | C+1- | 1.000 |
| | | 60 | 3.2 | | | 0.995 |
| | | 90 | 2.6 | | | 0.985 |
| | | 180 | 1.85 | | | 0.998 |
| | Right-STN | 140 | 2.8 | | C+5- | 1.000 |
| | | 60 | 4.25 | | | 0.987 |
| | | 90 | 3.5 | | | 1.004 |
| | | 180 | 2.45 | | | 0.984 |
| P05 | Left-STN | 130 | 2.25 | 70 | C+1- | 1.000 |
| | | 60 | 3.3 | | | 0.993 |
| | | 90 | 2.7 | | | 0.997 |
| | | 180 | 1.9 | | | 0.987 |

| | | | | | | |
|---|---|---|---|---|---|---|
| | Right-STN | 130 | 2.3 | | C+5- | 1.000 |
| | | 60 | 3.4 | | | 1.009 |
| | | 90 | 2.75 | | | 0.990 |
| | | 180 | 1.95 | | | 0.995 |
| P06 | Left-STN | 130 | 1.6 | 70 | C+3- | 1.000 |
| | | 60 | 2.35 | | | 0.996 |
| | | 90 | 1.9 | | | 0.976 |
| | | 180 | 1.35 | | | 0.986 |
| | Right-STN | 130 | 1.45 | | C+7- | 1.000 |
| | | 60 | 2.15 | | | 1.015 |
| | | 90 | 1.75 | | | 1.008 |
| | | 180 | 1.25 | | | 1.029 |

**Supplementary Table 2. Speech paradigms and feature definitions.**

| Speech task | Paradigm | Speech feature | Feature definition |
|---|---|---|---|
| Sustained vowel phonation | Sustain the vowel /a/ in a single breath. | HNR (dB) | Harmonics-to-noise ratio, defined as the ratio of harmonic components to noise components. |
| | | Jitter (%) | The average absolute difference between consecutive periods, divided by the average period. |
| | | Shimmer (%) | The average absolute difference between the amplitudes of consecutive periods, divided by the average amplitude. |
| Diadochokinetic (DDK) task | Repeatedly produce the syllable sequence /pa-ta-ka/ in a single breath. | DDK rate (syll./s) | Number of syllables produced per second. |
| Passage reading | Read aloud a 106-character Chinese passage at a comfortable speaking rate. The passage was designed to cover a wide range of Mandarin initials and finals with an approximately balanced distribution of lexical tones. | F0 Mean (reading) (Hz) | The mean fundamental frequency (F0) measured during the reading segment. |
| | | F0 Std (reading) (Hz) | The standard deviation of F0 measured during the reading segment. |
| | | F0 CV (reading) (%) | The coefficient of variation of F0 measured during the reading segment, calculated as F0 Std/F0 Mean. |
| Monologue | Speak freely for one minute on any topic in a natural speaking style, without reciting, chanting, or singing. | F0 Mean (monologue) (Hz) | The mean F0 measured during the monologue segment. |
| | | F0 Std (monologue) (Hz) | The standard deviation of F0 measured during the monologue segment. |
| | | F0 CV (monologue) (%) | The coefficient of variation of F0 measured during the monologue segment. |
| | | Speech Rate (char./s) | Number of characters produced per second. |
